# Functional Parcellation of the Default Mode Network: A Large-Scale Meta-Analysis

**DOI:** 10.1101/225375

**Authors:** Shaoming Wang, Lindsey J. Tepfer, Adrienne A. Taren, David V. Smith

## Abstract

The default mode network (DMN) consists of several regions that selectively interact to support distinct domains of cognition. Of the various sites that partake in DMN function, the posterior cingulate cortex (PCC), temporal parietal junction (TPJ), and medial prefrontal cortex (MPFC) are frequently identified as key contributors. Yet, it remains unclear whether these subcomponents of the DMN make unique contributions to specific cognitive processes and health conditions. To address this issue, we applied a meta-analytic parcellation approach used in prior work. This approach used the Neurosynth database and classification methods to quantify the association between PCC, TPJ, and MPFC activation and specific topics related to cognition and health (e.g., decision making and smoking). Our analyses replicated prior observations that the PCC, TPJ, and MPFC collectively support multiple cognitive functions such as decision making, memory, and awareness. To gain insight into the functional organization of each region, we parceled each region based on its coactivation pattern with the rest of the brain. This analysis indicated that each region could be further subdivided into functionally distinct subcomponents. Taken together, we further delineate DMN function by demonstrating the relative strengths of association among subcomponents across a range of cognitive processes and health conditions. A continued attentiveness to the specialization within the DMN allows future work to consider the nuances in sub-regional contributions necessary for healthy cognition, as well as create the potential for more targeted treatment protocols in various health conditions.

## Introduction

The default mode network (DMN) underlies a striking array of divergent processes, from interpreting a social exchange to contemplating future plans. As a result of this broad functional spectrum, the DMN is a focal point of active interrogation: decades of research detail its involvement in a wide range of healthy cognitive functions and psychiatric illnesses^1–3^. For instance, early in its discovery, the DMN earned a reputation for self-referential processing^4^ and mind-wandering^5^ after evidence emerged that the network played a role in spontaneous thought processes when study participants were not actively engaged in a directed task. Later, studies involving autobiographical memory retrieval^6^, self-judgements^7,8^, prospective thinking^6^, decision-making^9,10^, and social cognition^11^ were also found to evoke activation across the DMN, indicating that the same networked areas may play an indirect role across these psychological processes^10^. Ongoing research revealed that the DMN tended to engage many of the same subregions, particularly the medial prefrontal cortex (MPFC), posterior cingulate cortex (PCC), and left and right temporal-parietal junction (left- and right-TPJ)^12,13^. These four regions have been shown to preferentially activate when the brain is at rest, and decrease when engaged in a goal-directed task^14^.

In addition to studies linking the DMN to various cognitive processes, recent efforts have explored the DMN’s role in health problems, including psychopathology and neurological disorders. A growing body of literature suggests that DMN dysfunction may underlie disease states including Alzheimer’s disease, schizophrenia, ADHD, Parkinson’s disease, depression, and anxiety^15^. Decreased activity of the DMN at rest and decreased task-induced deactivation of the DMN has been observed in individuals with autism^16,17^, particularly in the MPFC^18,19^. Patients with anxiety disorders show reduced deactivation of MPFC and increased deactivation of PCC^15^, while the component regions of DMN appear to change during major depressive episodes, with activity of thalamus and subgenual cingulate increasingly seen at rest^20^. Alzheimer’s patients not only show altered DMN responses at rest, but different task-induced deactivation patterns during a working memory task^21^ and regional activation differences within the PCC^22^. In schizophrenia, both resting state and task-based DMN response changes have been associated with positive disease symptoms^3,15,23^.

While the aforementioned disease states share the commonality of generally altered DMN function, the findings from these studies also suggest that altered activity in different DMN nodes may be specific to different health conditions; for instance, with the PCC specifically implicated in Alzheimer’s and ADHD, and the MPFC in schizophrenia and anxiety^15^. However, despite the growing body of knowledge regarding the roles of these four DMN subregions in both the cognitive and psychiatric domains, it remains unclear whether further subdividing these regions would reveal distinct patterns of activity that show relative differences among one another in their contributions to various cognitive process and mental disorders alike.

Evidence for subregional specialization for a range of cognitive functions has been heavily documented in the MPFC in particular, exemplifying how a region may play different roles in cognitive function dependent upon subregional activation^24–26^. For instance, the MPFC as a whole appears to play an important role in self-referential processing, but shows a divergence in activity among its subregions: the ventral MPFC deactivates more during self-referential judgments while the dorsal MPFC activity increases^8^. This ventral-dorsal subspecialization within the MPFC appears to extend to other cognitive domains, with ventral MPFC responsible for emotional processing and dorsal for cognitive function^8^. Recent meta-analysis further delineated regional differences in MPFC’s function, identifying anterior, middle, and posterior zones, responsible for episodic memory and social processing, cognitive control, and motor function, respectively^27^. This previous work suggests there may be overlap in function and sub-specialization within MPFC regions ^27^, opening the door for further research into the more fine-grained aspects of MPFC’s function.

Although these previous lines of investigation into DMN functional specialization have largely focused on the MPFC, the PCC and the bilateral TPJ may also serve a wide range of functions. For example, the PCC is thought to play a key role in focused attention during task-based activities^28,29^ and continuous monitoring of internal and external stimuli at rest^2^. It has also been implicated in retrieval of episodic memory^30,31^, emotional processing^32^, and self-referential processing^33^. Similarly, the TPJ^34^ has been shown to play a role in self-referential processing^4^, and is important for social cognition along with other DMN regions (PCC and MPFC^11,35^). Less is known about potential functional subspecialization within the PCC and TPJ, and it is possible that these DMN regions may also show intra-regional differences in cognitive processing similar to MPFC^36,37^.

To build a better understanding of whether cognitive processes and health conditions are linked to activation within distinct subunits of the DMN, we used the Neurosynth database and identified core DMN regions based on prior work and anatomical landmarks. We also used tools and methods developed in prior work^38,39^ that have successfully characterized the functional substructure of the medial prefrontal cortex^27^ and the lateral prefrontal cortex^40^. Specifically, we employed a naïve Bayesian classifier to quantify the association between activation in four particular DMN nodes—the PCC, TPJ, and MPFC activation—and topics pertaining to cognition and health. Our application of these analytical methods to four key regions of the DMN is important because it allowed us to quantify how those DMN regions are functionally associated with cognitive processes and disease states—thus extending and complementing prior efforts to fractionate the DMN^12,36,37,39,41–46^. Our primary analyses focused on two key questions. First, are different psychological functions and disease states preferentially associated with distinct nodes of the DMN? Second, are there functionally distinct subregions within individual DMN nodes?

## Results

To determine how the PCC, MPFC, and TPJ coactivated with the rest of the brain, we begun by searching for the term “DMN” in Neurosynth to create a functional mask, which we then constrained to the 4 anatomical regions of interest to the present study (PCC, MPFC, left and right TPJ) using the Harvard-Oxford probabilistic atlas in FSL (Fig 1A). Note that our TPJ region corresponded to what was labeled as “posterior-TPJ” in a previous study^47^. After generating the DMN mask through functional mapping, we applied k-means clustering to determine whether the four subregions of the DMN showed functional segregation. When evaluating coactivation and functional preference profiles for the whole DMN, our contrasts reflect comparisons across DMN nodes; when investigating functional segregation of each DMN node, contrasts within a single DMN region represent comparisons among the clusters within an individual region. We note that our instantiation of our naïve Bayesian classifier used to determine whether topics predicted activity in a region or cluster used a fourfold cross-validation scheme, and classification accuracy was assessed by calculating the mean score across all four folds, scored using the area under the curve of the receiver operating characteristic (AUC-ROC). Permutation-based analyses accounted for the variability in the standard deviation values across folds.

**Fig 1.**
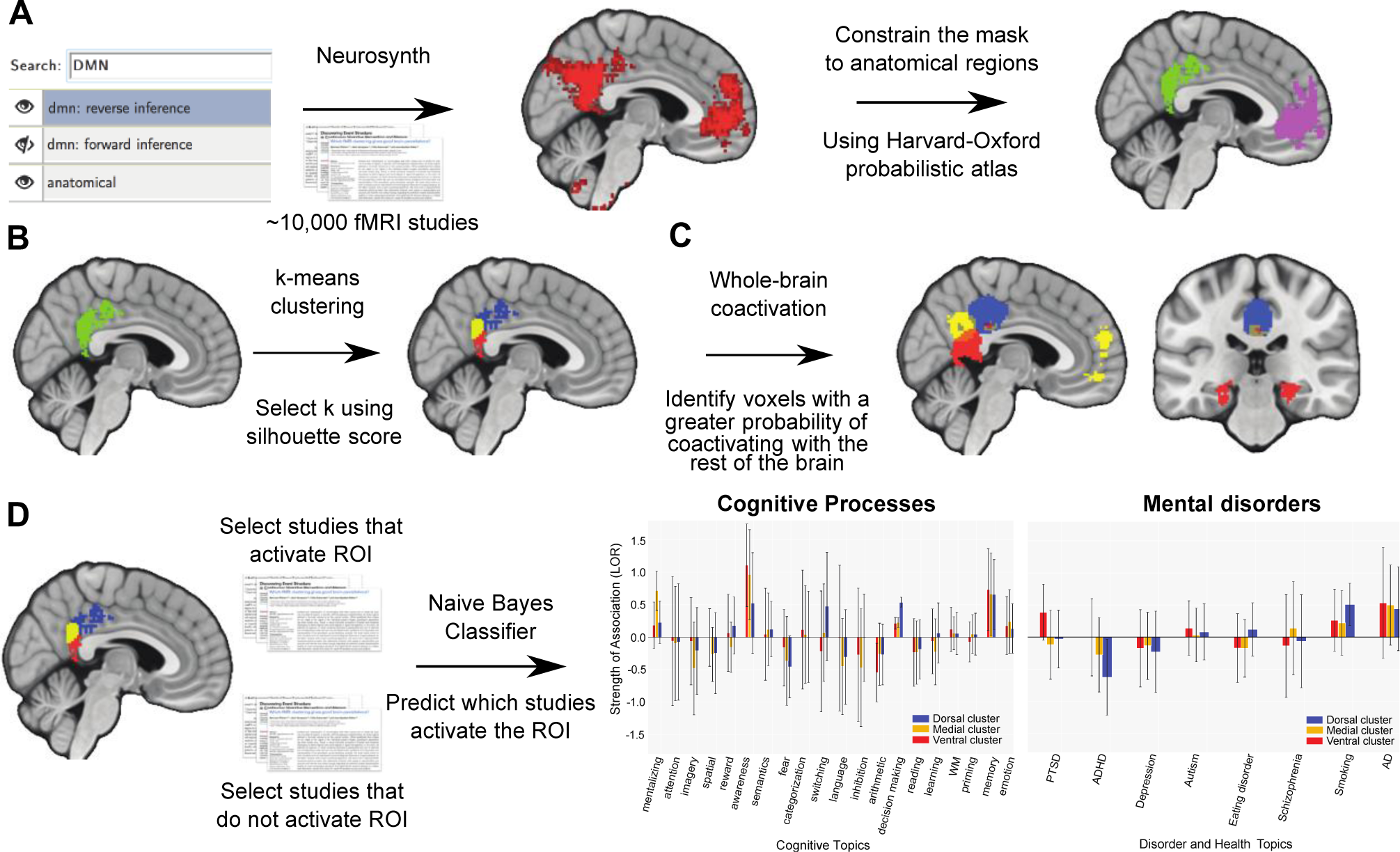
Overview of methods. (A) We searched for the topic “DMN” in Neurosynth to create a functional mask, and then constrained the mask to 4 anatomical regions that belong to the DMN by using the Harvard-Oxford probabilistic atlas. (B) We applied k-means clustering to determine functionally different subregions within each of the 4 regions. (C) We generated whole-brain coactivation profiles to reveal functional networks for different regions. (D) Functional profiles were generated to identify which cognitive processes or health conditions best predicted each region’s (or subregion’s) activity.

### Coactivation and functional preference profiles for the DMN regions

Next, we sought to characterize functional similarities and differences across regions of DMN. To determine how these four regions within the DMN coactivate with voxels across the brain, we adopted an approach used by previous work^27,40^, and identified voxels with a greater probability of coactivating with each ROI relative to the other regions within the DMN. First, we found that the four ROIs within the DMN coactivate with each other: PCC and MPFC strongly coactivated, and also showed coactivation with both the left and right TPJ (Fig. B). This pattern corresponds with long-standing evidence that coactivation among the various components within the DMN represents a typical characteristic of the network. In addition to the “collective” coactivation pattern among the four ROIs, we also found that the four regions showed coactivation overlap in ways that were not shared among all four ROIs. For instance, relative to the MPFC, the PCC and bilateral TPJ showed stronger coactivation with the hippocampus. Additionally, we found that relative to the PCC and bilateral TPJ, the MPFC showed stronger coactivation with both the amygdala and the ventral striatum, regions known for emotion processing and decision-making (Fig. B). Putting our coactivation results in perspective, these patterns demonstrate that there are both functional similarities and differences that characterize how these four regions within the DMN may operate relative to one another.

To further explore functional properties among DMN regions, we performed a meta-analysis to select studies that activated a given ROI, and then trained a naive Bayesian classifier to predict which studies activated the region^27^. From our classifier, we extracted the log odds ratio to determine whether a topic predicts activation in a particular cluster, with positive values indicating that a topic is indeed predictive of cluster activation. We first described functional preference profiles based on cognitive processes, followed by those related to health conditions. Cognitive predictors were limited to 20 psychological topics— words that represent co-occurrence among fMRI study abstracts within the Neurosynth database, e.g., cognitive processes and health conditions—previously shown to be relevant to DMN function. We found that all four regions of interest within the DMN were primarily associated with the topics “social”, “decision making”, “awareness”, and “memory” (Fig. 1C), consistent with previous evidence suggesting DMN involvement across these domains. We also found that activity among the four DMN regions were differentially associated with certain topics: fear, emotion and reward topics were associated with activity in the MPFC; emotion topics were associated with PCC activity; and math, semantics, and reading topics were associated with activity in the left TPJ. Next, we assessed the 8 disorder-related topics to examine whether regions within the DMN were differentially associated with various health conditions as well. We found that some disorders were associated with the MPFC or PCC: topics relating to smoking, eating disorder and depression were associated with activity in the MPFC, and topics pertaining to smoking and Alzheimer’s/Parkinson’s disease were associated with activity in the PCC (Fig. 1D). These results are consistent with observed coactivation patterns among regions within the DMN, again supporting the notion that there are functional similarities as well as differences among these four regions of the DMN, including function relating to psychiatric and health conditions.

Next, we evaluated whether the strengths of associations between a topic and a particular region were significantly distinct compared to other regions within the DMN. Our post hoc analysis suggested that despite their associations, none of the topics showed a statistically greater preference for one region over another within the DMN (or any of the subregions in the results that follow). Note that we caution interpretation of these results because these comparisons were post hoc and exploratory ^27^.

### Functional Distinctions Within DMN Subregions

We clustered individual voxels inside the MPFC, bilateral-TPJ and PCC based on their meta-analytic coactivation with voxels in the rest of the brain to elucidate more fine-grained functional differences among subregions^27,48,49^. We used silhouette scoring to select optimal solutions for each region, and generated coactivation and functional preferences profiles for each subregion. We describe the results for four regions separately.

Within the MPFC, we identified two clusters based on the silhouette score (Fig. 3A, left panel): a dorsal cluster (Fig. 3A right panel, orange) and a ventral cluster (Fig. 3A right panel, green). Our analysis did not reveal that either cluster coactivated more strongly with the rest of the brain. When exploring which topics predicted activation in these clusters, we found that both the dorsal and ventral MPFC clusters were associated with social, emotion, reward, decision-making, awareness, and memory; whereas the ventral cluster was associated with fear (Fig. 3B). Additionally, both clusters in MPFC were associated with depression and smoking, and the ventral cluster was associated with eating disorders (Fig. 3C; again, with no significant difference between the strength of association among cluster and topic associations).

**Fig 2.**
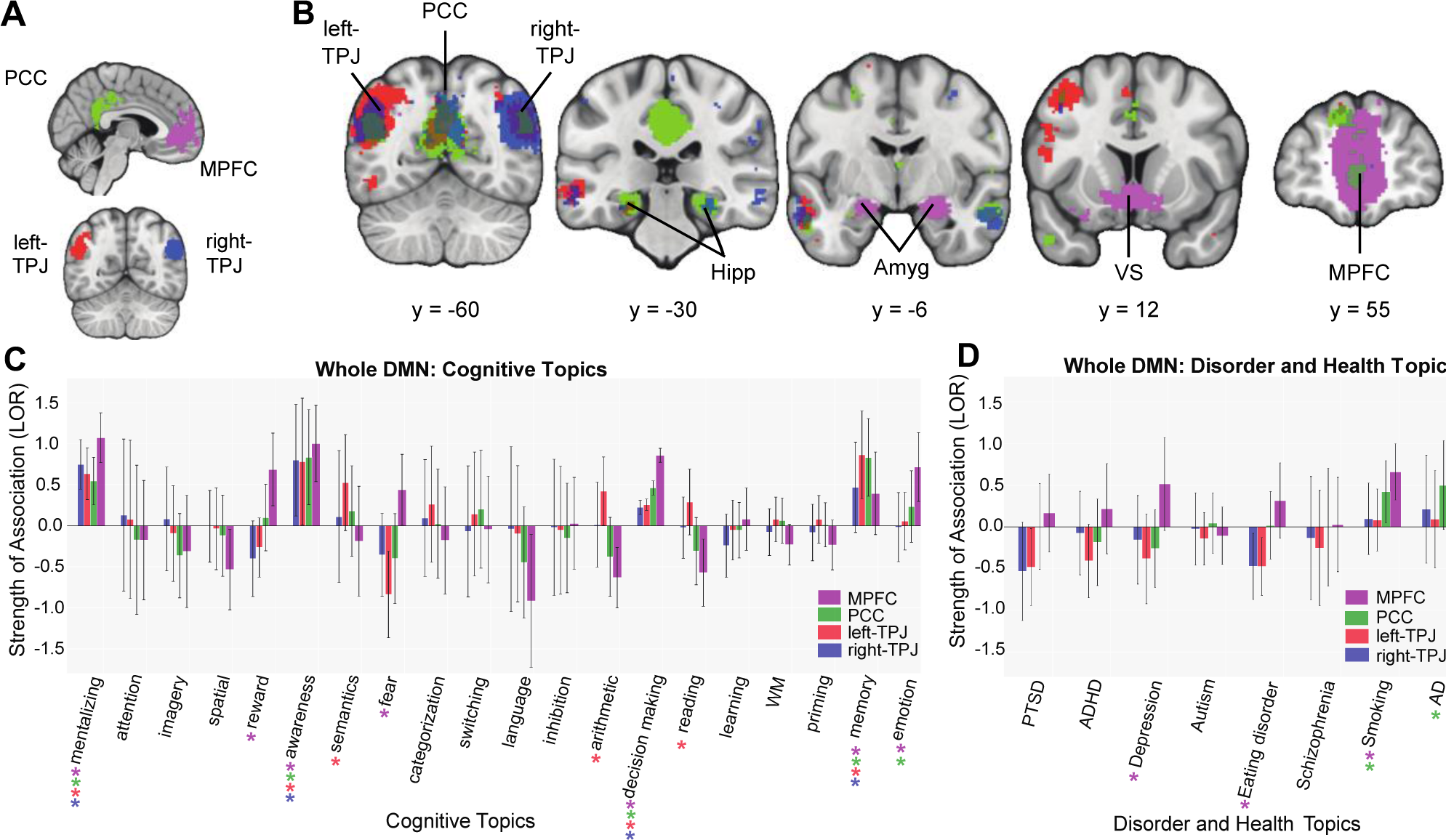
Meta-analytic coactivation and functional profiles for the DMN. (A) Functionally defined regions within DMN: medial prefrontal cortex (MPFC), posterior cingulate cortex (PCC), left and right temporal parietal junction (left- and right-TPJ). (B) Coactivation profiles for 4 regions. Colored voxels indicate significantly greater coactivation with the region of same color (Fig. 2A) than control regions. Some regions were involved in overlapping functional networks whereas some did not show the same degree of overlap across functional networks. Related subcortical structures are labeled as: Hipp, hippocampus; Amyg, amygdala; VS, ventral striatum. **Strength of association with cognitive and disorder topics measured by LOR.** (C) Functional profiles were generated by determining which cognitive topics best predicted each DMN cluster’s activity. All regions within DMN were primarily involved with social, decision making, awareness, and memory. Distinct functions were also observed across MPFC, PCC and left-TPJ. Log-odds ratio (LOR) is used to measure strength of association, and color-coded dots corresponding to each region are used to indicate significance (p < 0.05, corrected) based on permutation testing. (D) DMN and health conditions. Functional profiles related to health conditions were generated to determine whether regions within DMN were differentially recruited by psychological diseases.

**Fig 3.**
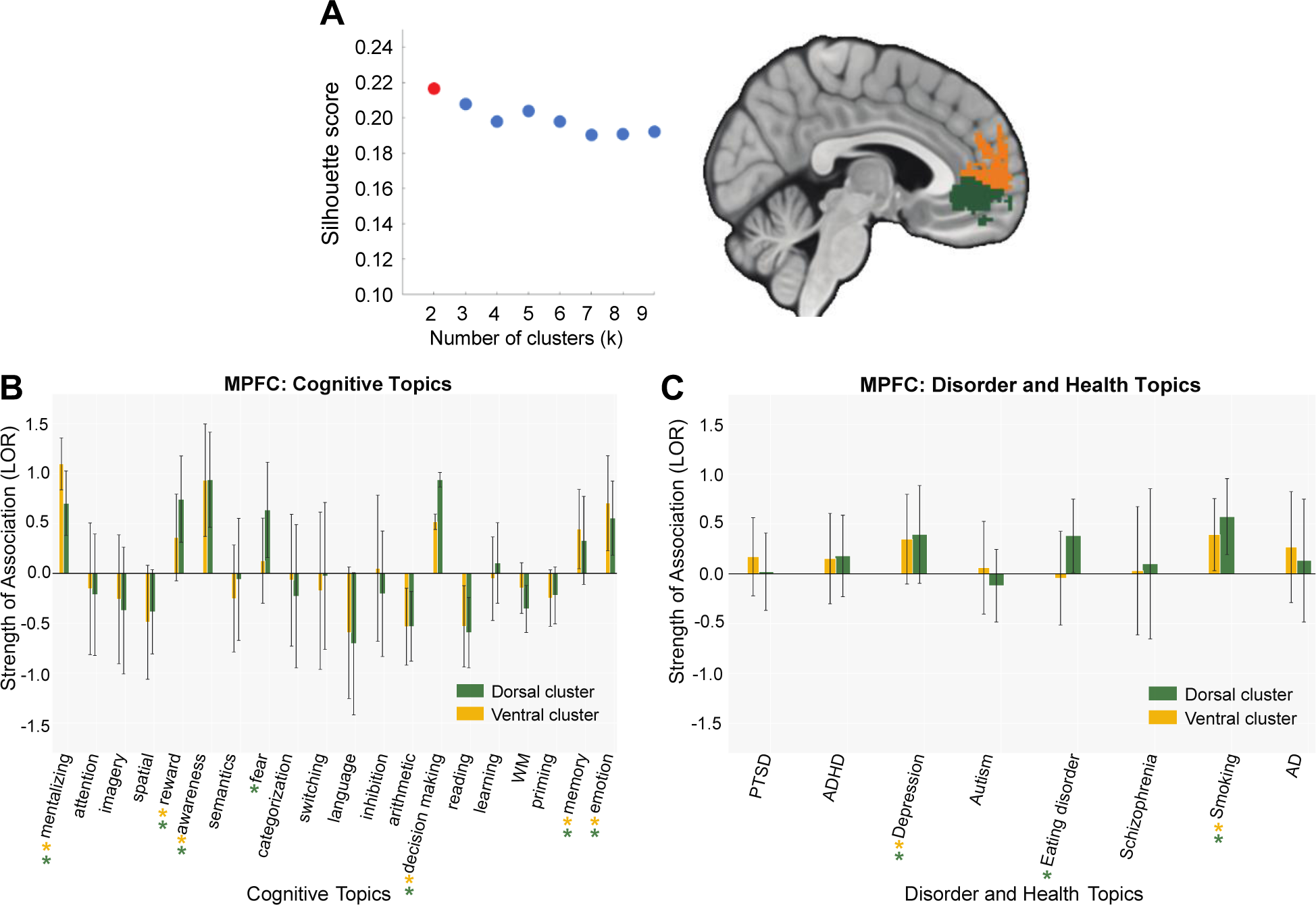
Coactivation-based clustering and functional preference profiles of MPFC. (A) We identified two clusters within MPFC based on silhouette scoring: a dorsal cluster (orange) and a ventral cluster (green). **Strength of association with cognitive and disorder topics measured by LOR.** (B) Functional preference profiles of the MPFC for cognitive topics. Both clusters in MPFC were predicted by social, emotion, reward, decision-making, awareness and memory, whereas the ventral cluster was predicted by fear. (C) MPFC and health conditions. Both clusters in MPFC were recruited by depression and smoking, but only the ventral cluster was associated with eating disorders.

Our silhouette score analysis revealed that a three-cluster solution was optimal for the right-TPJ (Fig. 4A, left panel): a dorsal cluster (Fig. 4A, green), a ventral one (Fig. 4A, yellow) and a posterior cluster (Fig. 4A, purple). All three clusters were strongly coactivated with left-TPJ (Fig. 4B). The ventral cluster showed more coactivation with the PCC. Although topic-cluster associations did not differ amongst each other, activity in all three clusters were associated with social, memory, and awareness topics, with decision making associated with the dorsal cluster (Fig. 4C). No subregions showed any association with disorder-related topics (Fig. 4D).

**Fig 4.**
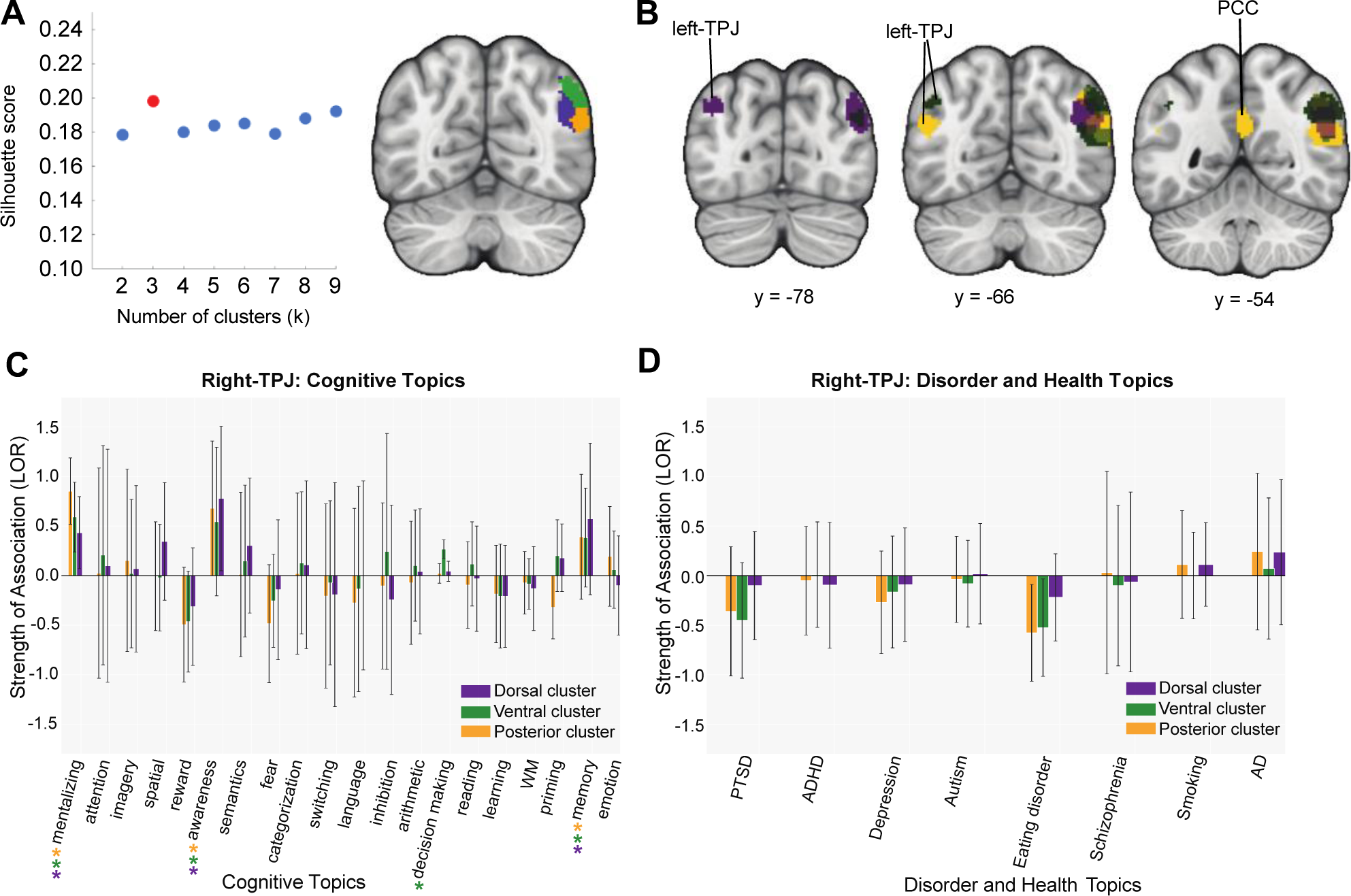
Coactivation-based clustering and functional preference profiles of right-TPJ. (A) We identified three clusters within the right-TPJ using silhouette scoring: a dorsal cluster (green), a ventral cluster (yellow) and a posterior cluster (purple). **Strength of association with cognitive and disorder topics measured by LOR.** (B) Coactivation contrasts for right-TPJ. All clusters strongly coactivated with left-TPJ. The ventral cluster showed more coactivation with PCC. (C) Functional preference profiles of right-TPJ. All clusters in right-TPJ were predicted by social, memory, and awareness, while only the dorsal cluster was predicted by decision making. (D) Right-TPJ and health conditions. No subregions were significantly associated with disease-related topics.

We identified three clusters within left-TPJ with silhouette scoring (Fig. 5A, left panel): an anterior cluster (Fig. 5A right panel, red), a posterior cluster (Fig 5A right panel, yellow) and a ventral cluster (Fig. 5A right panel, green). We directly contrasted the coactivation patterns of each subregion. We found all three regions strongly coactivate with right-TPJ. In addition, the posterior and ventral clusters showed stronger coactivation with PCC (Fig. 5B). While both clusters also strongly coactivated with MPFC, the posterior cluster showed strong coactivation with ventral MPFC, whereas the ventral cluster coactivated with dorsal MPFC more (Fig 5B). Consistent with functional patterns typically associated with the DMN, all clusters in the left-TPJ were primarily predicted by social, decision-making, memory, and awareness (Fig 5C). We found that the ventral cluster demonstrated an association with reading, semantics and emotion; the anterior cluster showed an association with reading and working memory; and the posterior cluster was associated with priming (Fig. 5C). No subregions within the left-TPJ showed any association with disorder-related topics (Fig. 5D). Note that the parcellations within left- and right-TPJ partially mirror each other (Table 1), which increases confidence in the clustering of coactivation patterns ^27,40^.

**Table 1.**
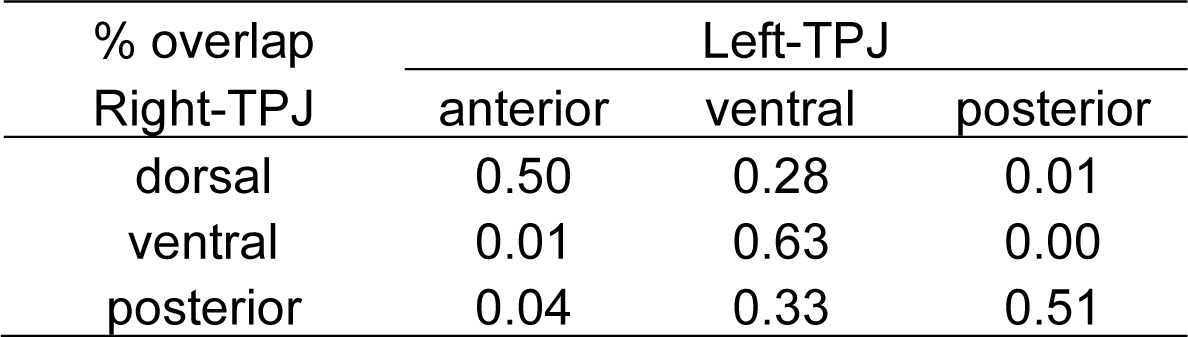
Percent overlapping voxels within subregions of left- and right-TPJ.

| % overlap<br>Right-TPJ | Left-TPJ |  |  |
| --- | --- | --- | --- |
|  | anterior | ventral | posterior |
| dorsal | 0.50 | 0.28 | 0.01 |
| ventral | 0.01 | 0.63 | 0.00 |
| posterior | 0.04 | 0.33 | 0.51 |

**Fig 5.**
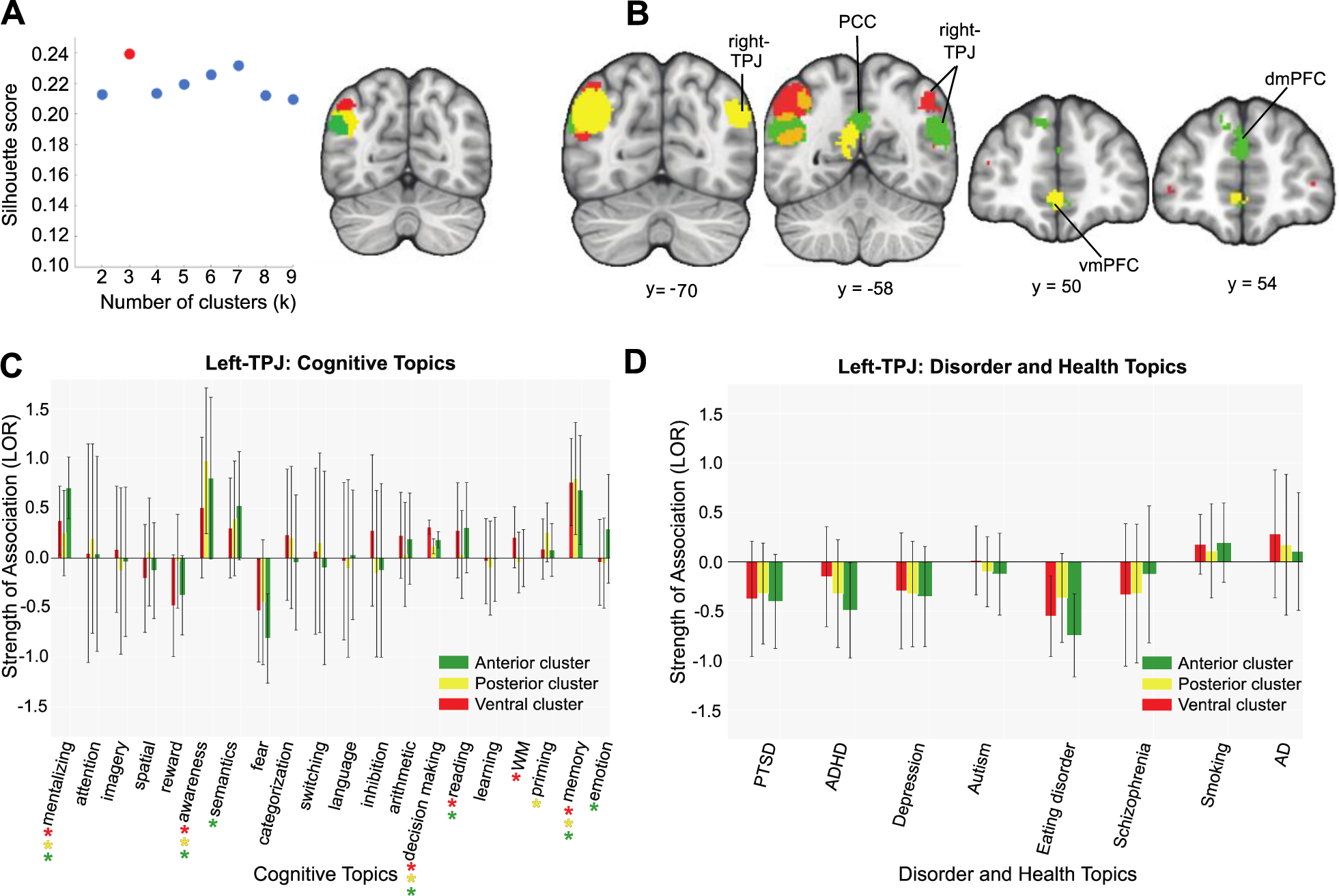
Coactivation-based clustering, meta-analytic coactivation contrasts and functional preference profiles of left-TPJ. (A) We identified three clusters within the left-TPJ based on silhouette scoring: an anterior cluster (red), a posterior cluster (yellow) and a ventral cluster (green). (B) Coactivation contrasts of left-TPJ. All three regions strongly coactivate with right-TPJ. The posterior and ventral clusters showed stronger coactivation with the right-TPJ. The posterior cluster showed more coactivation with the ventral MPFC whereas the ventral cluster more strongly coactivated with dorsal MPFC**. Strength of association with cognitive and disorder topics, of left-TPJ measured by LOR (Functional preference profiles).** (C) All clusters in the left-TPJ were primarily predicted by social, decision-making, memory, and awareness. The ventral cluster was more strongly associated with reading, semantics and emotion whereas the anterior cluster showed stronger association with reading and working memory. The posterior cluster was more strongly predicted by priming. (D) Left-TPJ and health conditions. No subregions showed strong associations with disease-related topics.

We identified three clusters within PCC with silhouette scoring (Fig. 6A, left panel): a dorsal cluster (Fig. 6A right panel, blue), a medial cluster (Fig. 6A right panel, yellow) and a ventral cluster (Fig. 6A right panel, red). The medial cluster showed stronger coactivation with bilateral-TPJ and MPFC, whereas the ventral cluster more strongly coactivated with hippocampus (Fig. 6B). Similar to this coactivation pattern, all clusters in PCC were predicted by memory, awareness, and decision-making, with the medial region showing an association with social cognition and emotion topics (Fig. 6C). When looking at the disorder topics, we found that all PCC clusters were associated with Alzheimer’s/Parkinson’s disease; the dorsal cluster was associated with smoking; and the ventral cluster was associated with PTSD (Fig. 6D), although these cluster-topic associations between the dorsal and ventral clusters were not significantly different in strength compared to each other.

**Fig 6.**
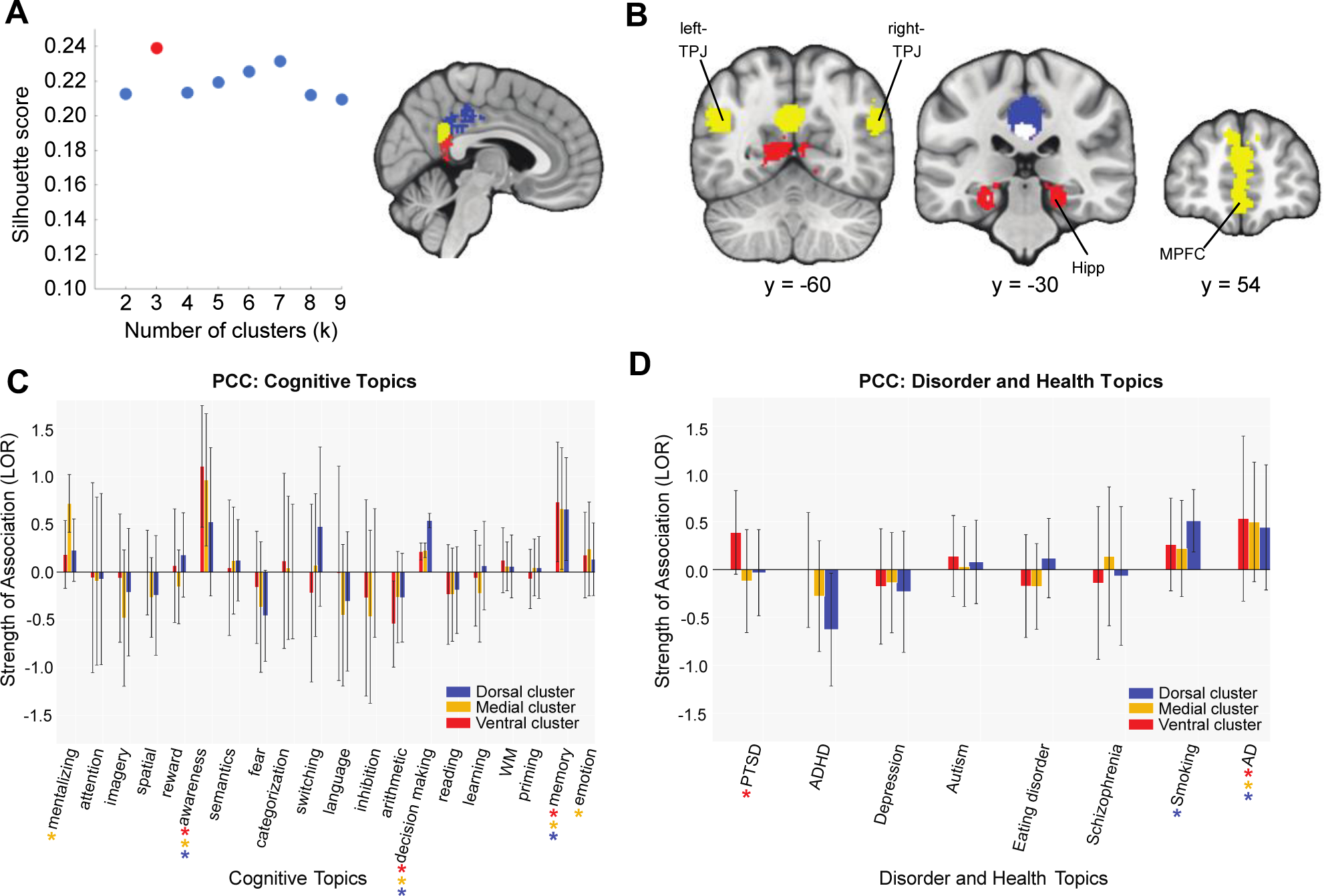
Coactivation-based clustering, meta-analytic coactivation contrasts and functional preference profiles of PCC. (A) We identified three clusters within the PCC on the basis of silhouette scoring: a dorsal cluster (blue), a medial cluster (yellow) and a ventral cluster (red). (B) Coactivation contrasts of PCC. The medial cluster showed stronger coactivation with the bilateral-TPJ and MPFC whereas the ventral cluster more strongly coactivated with hippocampus. (C) Functional preference profiles of PCC. All clusters in the PCC were predicted by memory, awareness, and decision making, while only the medial region was associated with social and emotion. (D) PCC and health conditions. All PCC clusters were associated with Alzheimer’s/Parkinson’s disease. The dorsal cluster was more strongly associated with smoking whereas the ventral cluster was more associated with PTSD.

## Discussion

Previous research has linked the default mode network to a wide range of cognitive functions and disease states. Numerous studies have elucidated the functional architecture of the network through coactivation- or connectivity-based analysis^36–39,42,45,50,51^. Yet, a comprehensive understanding of the mappings between psychological functions and different nodes within the default mode network remains in its early stages. To address this question, the present study builds upon previous work that used a high-powered meta-analysis of functional neuroimaging data to parcellate the MPFC and LFC by topic modeling of psychological functions^27,40^. Our findings help further characterize the functional specialization and subspecialization in two other key DMN regions, the PCC and TPJ. Here, we report coactivation among DMN regions and between DMN and hippocampus, amygdala, and striatum. Overlap in function among DMN regions was observed with all four ROIs for social cognition, decision-making, awareness, and memory. DMN regions were also differentiated by function, with MPFC associated with emotion, reward, and fear; PCC associated with emotion; and left-TPJ associated with math, semantics, working memory, and reading. Moreover, we found that two of the DMN regions were associated with psychopathology and neurological diseases: we found that the PCC was associated with Alzheimer’s, Parkinson’s disease, and smoking and the MPFC was associated with smoking, eating disorders and depression. Further examination of these DMN ROIs revealed that they could be divided into subregions based on cognitive and disease-related functional subspecialization.

Across all four DMN nodes, we observed consistent patterns of involvement with social cognition, memory, decision making and awareness. Our analyses also showed strong coactivation patterns between PCC and TPJ, suggesting their interactive roles in support of functions, such as social cognition and memory processes ^36,37,47^. These findings recapitulate the notion that nodes within DMN jointly contribute to multiple psychological processes ^7,14,45^. In addition, our results also revealed that the MPFC, as compared to other nodes, has more heterogeneous functional characteristics. We show that the MPFC coactivates with subcortical regions, such as the amygdala and ventral striatum. Consistent with this pattern, our analysis further shows that the MPFC is strongly associated with fear, emotion and reward, consolidating the vast literature on MPFC’s interactions with regions related to affect and reward learning^11,52^.

Within the MPFC, our finding of a strong association between the ventral cluster and fear is consistent with prior work implicating vmPFC in extinction of conditioned fear^53,54^. Our analyses further delineate psychological disorders associated with the MPFC. In particular, depression, which has been previously associated with altered DMN patterns^55^, loaded most strongly onto the MPFC. In patients with major depressive disorder, increased functional connectivity of the MPFC has been observed^56^, as well as increased MPFC activation during self-referential processing^57^. The fact that both dorsal and ventral MPFC clusters were associated with depression lends further support to previous work suggesting involvement of both MPFC subregions in major depressive disorder, with dorsal MPFC activation during depressive self-comparisons and ventral MPFC activation during the attentional component of depressive self-focus^58^.

Consistent with previous literature on PCC, the present study shows associations between PCC and social cognition. It has been shown that PCC is a central node of DMN for social cognition – e.g., ascribing mental states to others^11,12,59^ and social feedback^60^— and has functional interactions at rest with inferior parietal and superior temporal regions^11^. While previously the role of PCC in social cognition has not been well-delineated, our results suggest that a medial cluster of PCC may be most strongly associated with social cognition^61^. The same medial PCC cluster was shown to be specific to emotion; while there is relatively less evidence for the role of PCC in emotion, previous work has shown that PCC activates in response to emotional words and may mediate interactions between emotional and memory processes^32^, consistent with our parallel finding of an association between PCC and memory^62^. In addition to a primary association with memory, all PCC clusters loaded on Alzheimer’s/Parkinson’s disease, consistent with previous reports of PCC-DMN functional connectivity in patients with Alzheimer’s disease and mild cognitive impairment^63^. This association may also be linked to our finding of coactivation between PCC and hippocampus, as previous research has suggested disrupted resting state functional connectivity between PCC and hippocampus as a mechanism underling Alzheimer’s disease^63^.

Our analyses also highlight the association of both the dorsal and ventral PCC with smoking. Although most prior neuroimaging studies of smoking behavior focus on anterior cingulate and subcortical circuitry, nicotine has also been shown to consistently enhance functional connectivity between PCC and medial frontal/anterior cingulate cortex, as well as local connectivity between dorsal and ventral PCC^64^. Additionally, PCC activity has been associated with craving^65^, viewing smoking cessation messages^66,67^, and suppressing cue-induced craving^68^. Taken together, these functional distinctions indicate that subregions within PCC were differentially recruited by different cognitive processes, a pattern consistent with previous literature suggesting the multifaceted role of PCC in cognition^43,69,70^.

We also report involvement with social cognition within the TPJ, with the left TPJ more strongly associated with math, semantics, reading, emotion, and working memory. Within the domain of social cognition, previous research has established TPJ as most associated with mentalizing and theory of mind ^71^. In addition, previous work has suggested functional heterogeneity within TPJ on the basis of its functional and structural connectivity^11,41,72^. One study in particular attempted to map social cognition in the human brain, including parcellating TPJ using diffusion-weighted imaging with comparison to non-human primates^47^. This work suggested that posterior TPJ was most strongly associated with social cognition. Our analysis similarly shows strong loading of the ‘social’ term across all TPJ clusters; the previously reported social association with posterior and not anterior TPJ may be a result of TPJ mask definition, as Mars et al. (2011) included a broader anterior area of TPJ that overlaps further with the inferior parietal lobe.

Although our study provides a comprehensive characterization of the functional roles of the DMN, we note that our findings should be viewed with three caveats. First, the classifier used in our analysis did not distinguish activations from deactivations. However, it is well established that the DMN might be activated for some processes (e.g., social cognition^11,50,73^; and deactivated for others (e.g., executive control^74–76^). It is thus conceivable that a dataset capable of detecting deactivations would potentially extend our current findings and provide a full account of the functional architecture of the DMN. Second, the coactivation maps may not be directly related to connectivity between brain regions because they are based on correlations. Indeed, correlation between brain regions can be driven by a number of factors that are not related to connectivity or coupling, including changes in signal-to-noise ratio in either region or a change in another brain region^77,78^. A thorough examination of connectivity would necessitate integrating behavioral tasks with effective connectivity measures, such as psychophysiological interactions analysis^78–82^ (PPI), an avenue for future investigations. This alternative approach would provide insight into how specific tasks and processes drive connectivity with the DMN. Finally, the nature of the topic modeling oversimplifies the mapping between psychological ontology to complex, dynamic brain activity^40,83^. For example, each topic used in our analysis represents a combination of many cognitive processes operating at different levels. As a result, mappings between specific psychological concepts and brain activity may require further parcellation of complex cognitive processes.

In addition to these limitations, we also note that our approach for defining and fractionating the DMN merits additional consideration. For example, we defined the DMN based on a combination of anatomy and function and then parcellated individual nodes of DMN. Although an analogous approach has been used in prior studies^12^, we note that other studies have parcellated networks using responses from all nodes^61^. Both approaches assume a given network is composed of distinct nodes, which depends critically on the definition of those nodes^84,85^. To address this issue, some papers have defined networks using continuous maps (e.g., those derived from independent component analysis) and have examined connectivity with those maps using dual-regression analysis^86–88^. We believe that integrating this approach with our current analytical framework^27,40^ and unthresholded whole-brain maps^89^ will help future studies refine functional parcellations of the DMN. We also note that in our process of parcellating four regions within the DMN, our clusters had become relatively small, which may have limited our ability to find differences between cluster-topic associations among each DMN node. Future work may be able to address this limitation by using image-based meta analyses^90,91^, which may be possible as more researchers heed calls to share unthresholded statistical maps^82,89,92^.

To conclude, the present study applied a meta-analytic approach to characterize functional mappings among cognitive processes, health conditions, and the DMN. Although the DMN as a whole contributes to multiple cognitive processes, we identified distinct functional properties for each region, and extended these results by identifying further parcellation of functions for specific subregions. These results help clarify the functional roles of the DMN across a large corpus of neuroimaging studies and add to the body of literature that has focused on refining the theoretical and computational framework associated with the DMN^93^.

## Methods

Our analysis was based on version 0.6 of the Neurosynth dataset (i.e., articles up to July 2015), which contains activation data from more than 12,000 fMRI studies^94^.

### DMN mask

First, to find regions that are functionally related but restricted to the four anatomical regions within the DMN that we are interested in, we created a DMN mask that constrained wider functional activation within an anatomical atlas^95^ (Fig. 1A). Specifically, we performed a meta-analytical association test by searching the term “DMN” in Neurosynth to create a functional mask that specifically mapped the term “DMN” to brain regions. We chose an association test (formerly named a “reverse inference”) because it can help estimate the relative specificity between brain activity and the DMN ^94,96^. The resulting mask included 9,650 activations from 366 studies, and thus identified voxels in studies where the term “DMN” was mentioned in their abstract given brain activation. All of the images are thresholded with a false discovery rate of 0.01. Next, we constrained the mask to the four DMN regions of interest by using the Harvard-Oxford probabilistic atlas^97^ at P > 0.25, which included the medial prefrontal cortex (MPFC), the posterior cingulate cortex (PCC), and the left and right temporal parietal junction (left- and right-TPJ). This constraint was applied in the interest of specifying the output our functional mapping and subsequent parcellation to a precise set of standardized coordinates within the four regions of interest.

### Coactivation-based clustering

To determine more fine-grained functional differences within the four primary DMN regions of interest, we applied a clustering method used by previous work^27^ to cluster individual voxels inside each of the four regions based on their meta-analytic coactivation with voxels in the rest of the brain^38^ (Fig. 1B). To assess coactivation patterns for each of the four regions, we correlated the activation pattern of each voxel with the rest of the brain across all 366 studies. The resulting coactivation matrix is large and would require intensive computational power; to mitigate this, the coactivation matrix was passed through a principal component analysis (PCA), where the dimensionality of the matrix was reduced to 100 components^27,40^. Once the coactivation was reduced, we calculated the Pearson correlation distance between every voxel with each whole-brain PCA component in each of the four DMN regions. We then grouped the voxels within each separate region into 2-9 clusters by applying a k-means clustering algorithm to the correlation distance coefficients^98^. Finally, to determine the number of clusters for each region, we computed silhouette coefficients, allowing us to measure cohesion within each cluster^99^. Previous reports caution that issues may arise when identifying the appropriate number of clusters due to variations in goals and levels of analysis used across investigations ^83,100^; however, recent work has confirmed that the silhouette score can successfully assess cluster solutions^27,40,101,102^.

### Coactivation profiles

Next, we analyzed the how the whole-brain coactivation patterns differed between regions to reveal their functional networks^27^ (Fig. 1C). To accomplish this, we began by setting up contrasts among each of the four regions’ coactivation patterns (e.g., the MPFC against the PCC, left TPJ and right TPJ) to detect the coactivation differences between each region. Specifically, we performed a meta-analytic contrast between studies that activated the region of interest (ROI; “active studies”) and studies that activated the other three regions of interest (“inactive studies”), to identify the voxels in the rest of the brain that had a greater probability of coactivating with a given ROI than other regions within DMN. For instance, the purple voxels in Figure 2B indicates that these voxels in particular (in this case, within the amygdala and striatum) tend to co-activate alongside the MPFC rather than the PCC, the left TPJ or the right TPJ. To generate coactivation images that highlight coactivation profiles that emerged as a result of contrasting the four DMN ROIs, we conducted a two-way chi-square test between two sets of studies—the active and inactive studies—and calculated *p* values to threshold the coactivation images using False Discovery Rate (q<0.01). The resulting images were binarized and visualized using the NiLearn Library in Python ^103^.

### Meta-analytic functional preference profiles

To map between functional states (e.g., cognitive processes and health conditions) and the MPFC, PCC, and TPJ regions of the DMN, we re-purposed a set of 60 topics— generated as a part of a previous study^27^—which represent words that co-occur among fMRI study abstracts in the Neurosynth database. The set of 60 topics, which were produced in the exact same manner as seen in Poldrack et al., 2012, were derived from Dirichlet allocation topic modeling^104^, which is beneficial in reducing the level of redundancy and ambiguity in term-based meta-analysis maps in Neurosynth^27^. Per Poldrack et al., 2012 and De La Vega et al., 2016, topics that were irrelevant to either cognitive processes or health conditions (for instance, topics pertaining to experimental methods and design; N=23 topics) were excluded from the original set of 60 generated topics. After the exclusion step, we were left with 37 topics: 29 cognitive topics (e.g., mentalizing, attention, imagery) and 8 disorder-related topics (e.g., PTSD, ADHD, Autism).”

We generated functional preference profiles by identifying which cognitive processes or health conditions best predicted activity within a given cluster across studies (Fig. 1D). Considering the redundancy and ambiguity in term-based meta-analytic maps (e.g., “memory” could refer to working memory or episodic memory), we followed the lead of Yarkoni et al (2011), and trained a naive Bayesian classifier^27^ to evaluate two sets of studies. One set of studies included those that activated a cluster and the other set did not; a study was defined as activating a given cluster if at least 5% of voxels in the cluster were activated in that study. The classifier was then trained to predict whether a study activated a particular cluster, given the semantic representation of the latent conceptual structure underlying descriptive terms featured in the study^27,40^. Our instantiation of the classifier employed a fourfold cross-validation scheme, the default variance smoothing in scikit-learn (i.e., 1e-09), and did not utilize any priors. Input variables included the 60 psychological and health topics to predict whether activation is likely given the latent concepts within a study. As a final measure of our model’s performance, we calculated the mean score across all four folds, and scored them using a classification summary metric that considers the sensitivity and specificity: the area under the curve of the receiver operating characteristic (AUC-ROC).

Next, we extracted the log-odds ratio (LOR) of a topic, defined as the log of the ratio between the probability of a given topic in studies that activated a given cluster ROI and the probability of the topic in studies that did not activate the same cluster ROI, for each cluster separately. A positive LOR value indicates that a topic is predictive of activation in a given cluster. Based on LOR values, we identified 20 cognitive topics that loaded most strongly to the entire DMN mask for further analysis and then applied a procedure used in a previous study to determine the statistical significance of these associations^27^. To do so, we performed a permutation test for each region-topic log odds ratio 1000 times. This resulted in a null distribution of LOR for each topic and each region. We calculated *p* values for each pairwise relationship between topics and regions and then adjusted the *p*-values using a False Discovery Rate at 0.01 to account for multiple comparisons within 20 selected cognitive topics and 8 disease-related topics (separately). We reported associations significant at the corrected *p* < 0.05 threshold. Finally, to determine whether certain topics showed greater preference for one region or cluster versus another, we conducted exploratory, post hoc comparisons by determining whether the 99.9% confidence intervals (CI) of the LOR of a specific topic for one region or cluster overlapped with the 99.9% CI of the same topic for another region. We use a more stringent threshold than that of previous work^27^ due to the increased number of comparisons we make across DMN regions. We used bootstrapping to generate CIs, sampled with replacement, and recalculated log-odds ratios for each cluster 1000 times.

## Acknowledgments

We thank Alejandro de la Vega for sharing his analysis code and answering technical questions. We also thank Victoria Kelly and Christian Reice for their comments on an earlier draft of the manuscript. This work was supported in part by NIH grant R21-MH113917 (DVS).

## Data and code availability

All data are publicly available on Neurosynth database (http://neurosynth.org). In addition, all analysis code can be found on our lab GitHub (https://github.com/DVS-Lab/dmn-parcellation). All images can be found on NeuroVault (https://identifiers.org/neurovault.collection:6262).

## Author Contributions

S.W. participated in: formal analysis, writing - original draft, writing – review & editing, visualization. A.A.T. participated in: writing - original draft, writing – review & editing. L.J.T participated in: writing – review & editing, validation, visualization. D.V.S. participated in conceptualization, supervision, formal analysis, writing – original draft, writing – review & editing.

